# Vγ9+Vδ2+Tcell-mediated purging of *Listeria monocytogenes*-infected epithelial cells requires butyrophilin 3A genes

**DOI:** 10.1101/2023.06.07.544000

**Authors:** Katrin Fischer, Michaela Bradlerova, Thomas Decker, Verena Supper

## Abstract

Intracellular bacteria produce antigens, which serve as potent activators of γδ T cells. Phosphoantigens are presented via a complex of Butyrophilins (BTN) to signal infection to human Vγ9+Vδ2+ T cells. Here, we established an in vitro system allowing for studies of Vγ9+Vδ2+ T cell activity in coculture with epithelial cells infected with the intracellular bacterial pathogen *Listeria monocytogenes*. We report that the Vγ9+Vδ2+ T cells efficiently purge such cultures from infected cells. This effector function requires the expression of members of the BTN3A family on epithelial cells. Specifically, the BTN3A1 and BTN3A3 are redundant in their ability to present antigen to Vγ9+Vδ2+ T cells. Since BTN3A1 is the only BTN3A associated with phosphoantigen presentation our study suggests that BTN3A3 may present different classes of antigens to mediate Vγ9+Vδ2+ T cell effector function against L. monocytogenes-infected epithelia.

## 1. Introduction

The Gram-positive bacterium *Listeria* (*L*.) *monocytogenes* is causative agent of food-borne diseases. Infection via contaminated food may cause systemic spread of the bacteria and result in life-threatening meningoencephalitis or, in case of maternofetal infections, fetal abortion (reviewed in^1^).

Systemic bacterial spread results from *L. monocytogenes*’ ability to cross the intestinal, blood–brain and placental barriers. Virulence factors encoded by the *L. monocytogenes* pathogenicity islands endow the bacterium with the ability to force uptake into the cytoplasm of epithelial cells as well as intracellular replication and cell-to-cell spread. In phagocytic hosts *L. monocytogenes* escapes from phagosomes to replicate in the cytoplasm and spread between host cells (reviewed in^2,3^).

Defense mechanisms against intracellular bacterial infections include macrophage activation and the mobilization of cytotoxic T cells (reviewed in^4^). In humans, a special cytotoxic T cell subset of Vγ9+Vδ2+T cells recognizes bacterial antigens (reviewed in^5^). Vγ9+Vδ2+T cells expand in the peripheral blood in patients with *L. monocytogenes* infection, exceeding the levels of age-matched controls by 5-6-fold on average ^6^. The Vγ9+Vδ2+T cell expansion among peripheral blood mononuclear cells (PBMCs) could be recapitulated *in vitro* by exposing the cells to either viable *L. monocytogenes*^7^ or extracts from heat-killed bacteria^6,8^. Consistent with this, Vγ9+Vδ2+ T cell expansion was reproduced in non-human primate *L. monocytogenes* infection models^9,10^. Tanaka and colleagues identified (E)-4-Hydroxy-3-methyl-but-2-enyl Pyrophosphate (HMB-PP) as the immunostimulatory agent for the Vγ9+Vδ2+ T cell expansion. HMB-PP belongs to the group of phosphoantigens. It is a metabolite of the deoxyxylulose pathway for isoprenoid synthesis in *L. monocytogenes*^11^ and various other bacterial or parasitic pathogens^5^. Further support for the importance of HMB-PP was provided by extracts from bacterial mutants with increased HMB-PP levels that strongly stimulated Vγ9+Vδ2+T cell growth in a PBMC culture^12^ and from HMB-PP-deficient *L. monocytogenes* strains that revealed weaker Vγ9+Vδ2+ activating potential. The data underline a non-exclusive but important role of HMB-PP^13^.

BTN3A1 was identified and confirmed as important antigen-presenting molecule of HMB-PP and other phosphoantigens^14–17^. BTN3A is a member of the butyrophilins, plasma membrane-resident proteins related to the B7 costimulator family. The BTN3A family consists of 3 genes, BTN3A1, BTN3A2 and BTN3A3, however only BTN3A1 contains a positively charged pocket in its 30.2 domain, that was shown to bind phosphoantigens^18,19^. Very recently another butyrophilin family member, BTN2A1, was described to bind both BTN3A1 and the Vg9 T cell receptor (TCR) and to play an essential role in Vγ9+Vδ2+ T cell recognition^20–22^.

The majority of experimental approaches to study the impact of Vγ9+Vδ2+ T cells on *L. monocytogenes* infection is based on the use of bacterial extracts, synthetic phosphoantigens or drugs enhancing intracellular phosphoantigen levels. Noteworthy exceptions are studies in non-human primates^9^ and a report of Vγ9+Vδ2+ T cell cytotoxicity against infected monocyte-derived dendritic cells^5^. However, a system exploring the fate of *L. monocytogenes*-infected epithelial cells exposed to Vγ9+Vδ2+ T cells is missing. Epithelial cells represent the primary targets of gastrointestinal infection. The ability of host organisms to limit replication and spread of *L. monocytogenes* at epithelial barriers determines whether potentially life-threatening systemic infection will ensue.

The aim of this study was the establishment of an *in vitro* model to investigate consequences of subjecting infected epithelial cells to co-culture with Vγ9+Vδ2+ T cells. This system enabled us to monitor viability and the bacterial burden of target cells as well as measuring changes in the cytokine milieu. Gene deletion allowed us to scrutinize the roles of BTN3A genes. We observed redundant roles of BTN3A1 and BTN3A3 in allowing Vγ9+Vδ2+ T cells to purge cultures from infected cells, thus rescuing them from complete killing by *L. monocytogenes* infection. Changes of cytokine levels during co-culture largely depended on Vγ9+Vδ2+ T cells, *L. monocytogenes* infection and BTN3A expression. Our model extends the experimental options to address the role of Vγ9+Vδ2+ T cells in human infectious disease and identifies both BTN3A1 and BTN3A3 as antigen presenting molecules.^23^

## 2. Results

### 2.1. Vγ9+Vδ2+ T cells rescue cell populations from *L. monocytogenes*-induced killing

To fully understand the role of Vγ9+Vδ2+ T cells in *L. monocytogenes* infection, we established and optimized a co-culture system using the ‘real-time cell analysis’ (RTCA) method as a read-out. The RTCA system allows for continuous, label-free monitoring of cell adherence to study the time-resolved effects of Vγ9+Vδ2+ T cells on the growth and viability of infected target cells. Data are shown as % cell indices, where 100 represents the impedance (viability) of the culture at the onset of measurement. As a control for Vγ9+Vδ2+ T cell-induced cytotoxicity we used HDMAPP, a phosphoantigen and synthetic version of the bacterial Vγ9+Vδ2+ T cell activator HMB-PP. The detergent Triton X was used to mimic the effect of complete cell killing. As depicted in Figure 1A, Vγ9+Vδ2+ T cells alone caused an initial drop of viable RKO colon cancer cells, however, the cells recovered with progressive culture. In contrast, combination of both Vγ9+Vδ2+ T cells and the activator HDMAPP caused a dramatic drop in the cell index within the first 24 hours of co-culture and no subsequent recovery. This is in line with current literature showing a cytotoxic effect of Vγ9+Vδ2+ T cells on their target cells when treated with phosphoantigen^24,25^. In the *L. monocytogenes model*, about 5% of cells were infected with *L. monocytogenes*, at start of co-culture under the conditions used (Fig. S1). The cell index of *L. monocytogenes* infected target cells decreased steadily over 7 days as the infection was spreading from cell-to-cell. Strikingly, this drop was not observed in a co-culture system of Vγ9+Vδ2+ T cells and target cells infected with *L. monocytogenes*. Rather, a significant increase of the cell index was recorded, surpassing the cell index of *L. monocytogenes*-infected cells approximately 48 hours post infection. This Vγ9+Vδ2+ T cell-induced effect on infected target cells will henceforth be designated as the ‘rescue effect’. To determine whether the rescue effect is specific for RKO cells we performed RTCA assays with the ovarian cancer cell line COV362 (Figure 1B). Vγ9+Vδ2+ T cells exerted basal cytotoxicity on these cells, but a drop in cell viability in the presence of the phosphoantigen HDMAPP was similarly strong as in RKO cells. Upon *L. monocytogenes* infection, the cell index quickly dropped, and this effect was again rescued in co-culture with Vγ9+Vδ2+ T cells. To test a potential donor-specificity in the observed rescue effect, we performed the RTCA assay using a second Vγ9+Vδ2+ T cell donor and observed similar results (Figure 1C). In conclusion, Vγ9+Vδ2+ T cells rescue cell cultures from *L. monocytogenes*-induced cell death, thus enabling proliferation of the target cells over time. This effect is consistent across two different human epithelial cell lines as well as two independent Vγ9+Vδ2+ T cell donors.

**Figure 1.**
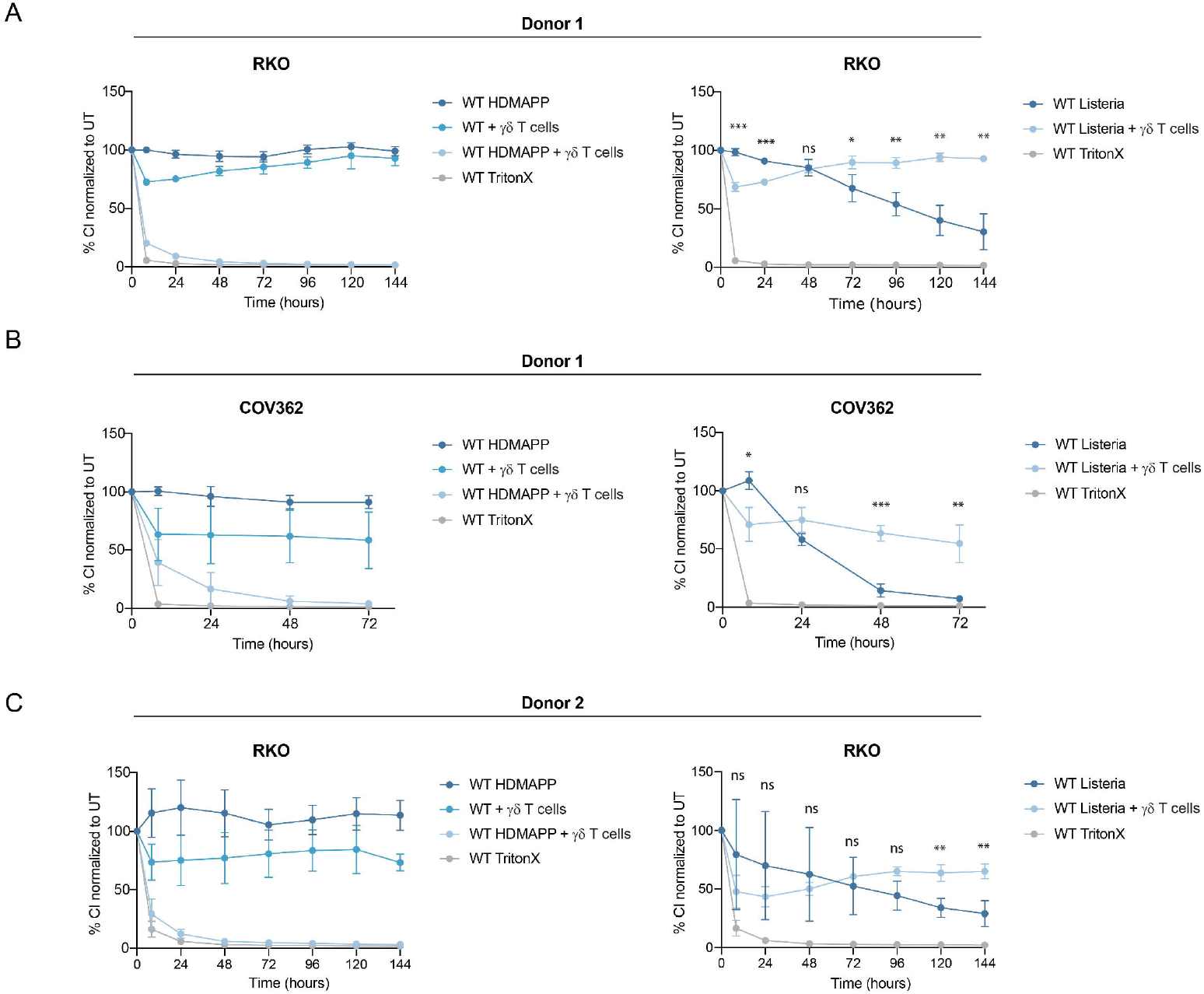
RTCA was performed with RKO **(A and C)** or COV362 **(B)** cells co-cultured with Vγ9+Vδ2+ T cells donor 1 **(A-B)** or donor 53X **(C)**. Target cells were either treated with HDMAPP (left panel) or infected with L. *monocytogenes (right panel)*. Triton-X was used as positive control for cytotoxicity. Cell Index of the treatments was normalized to the untreated sample (100%). Two-tailed unpaired Student’s t-test was performed using GraphPad Prism comparing L. *monocytogenes* with L. *monocytogenes* + Vγ9+Vδ2+ T cells. P-values: ns P > 0.05; *P ≤ 0.05; **P ≤ 0.01; ***P ≤ 0.001; ****P ≤ 0.0001. Experiments were performed in technical as well as biological triplicates.

### 2.2. The Vγ9+Vδ2+ T cell-mediated rescue of *L. monocytogenes*-infected target cells is BTN3A-dependent

Next, we addressed the question whether the observed rescue of *L. monocytogenes* infected cultures by Vγ9+Vδ2+ T cells requires members of the BTN3A family. Consistent with current literature^18,22,26,27^, BTN3A pan knockout cells lacking all three BTN3A genes (BTN3A1, BTN3A2 and BTN3A3) (Figure S2A), failed to succumb to the cytotoxic effect of Vγ9+Vδ2+ T cells in the presence of HDMAPP (Figure 2A). Likewise, *L. monocytogenes*-infected BTN3A pan knockout RKO cells co-cultured with Vγ9+Vδ2+ T cells did not exhibit the ‘rescue effect’ as their wildtype counterparts. The knockout cells experienced similar decreases of viability as *L. monocytogenes*-infected cells in absence of Vγ9+Vδ2+ T cells. The same results were obtained using BTN3A1/3 COV362 double-knockout cells (Figure 2B and Figure S2B). Thus, rescue of *L. monocytogenes*-induced cell death requires BNT3A1 or BTN3A3-dependent antigen presentation to Vγ9+Vδ2+ T cells in two human epithelial cell lines.

**Figure 2.**
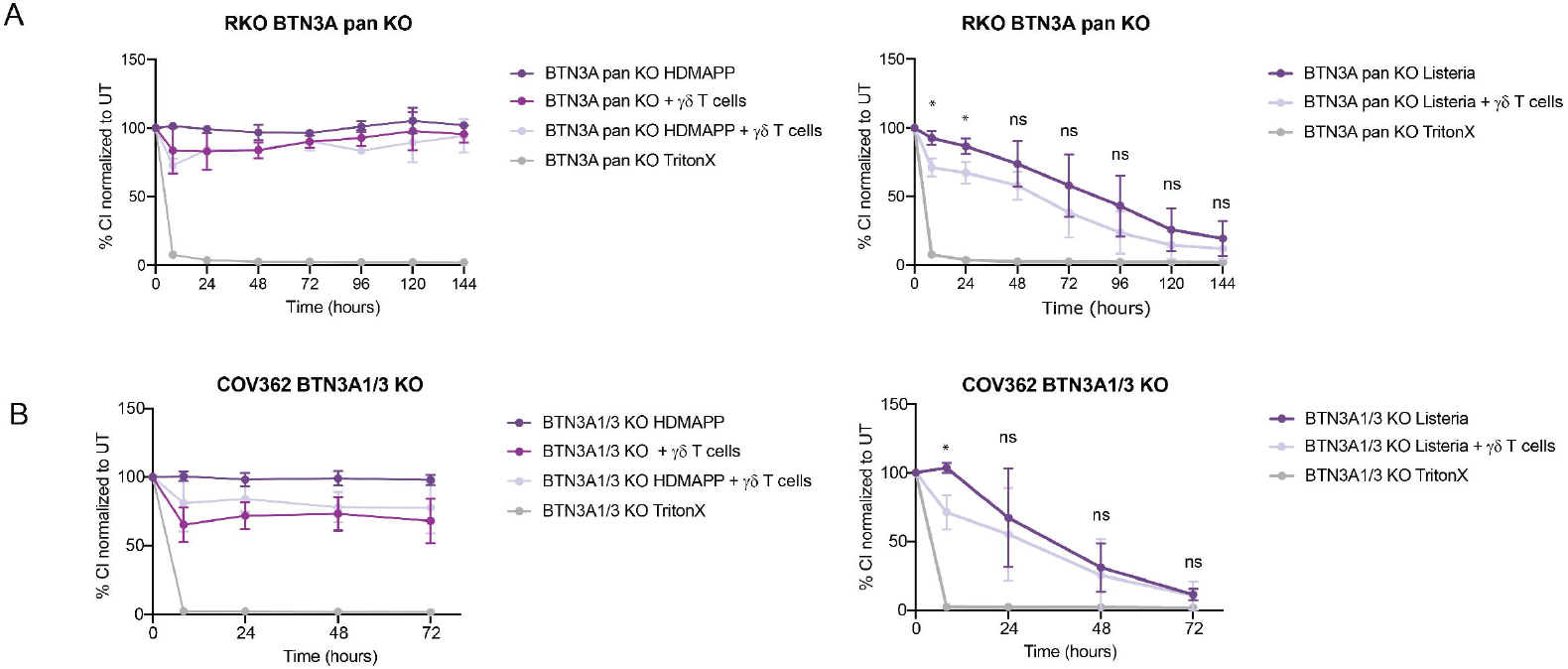
RTCA was performed with BTN3A pan knockout RKO **(A)** or BTN3A1/3 double knockout COV362 **(B)** cells in co-culture with Vγ9+Vδ2+ T cells from donor 1. TritonX was used as positive control for cytotoxicity. Target cells were either treated with HDMAPP (left panel) or infected with L. *monocytogenes* (right panel) Cell Index of the treatments was normalized to the untreated sample (100%). Two-tailed unpaired Student’s t -test was performed using GraphPad Prism comparing L. *monocytogenes* with L. *monocytogenes* + Vγ9+Vδ2+ T cells. P-values: ns P > 0.05; *P ≤ 0.05; **P ≤ 0.01; ***P ≤ 0.001; ****P ≤ 0.0001. Experiments were performed in technical as well as biological triplicates.

### 2.3. Cytokine production does not reveal clear correlations with BTN3A-dependent clearance of *L. monocytogenes* infection

Cytokines are key modulators of inflammation, shape the cellular environment during infections and reflect immune cell activation. In addition, they establish cell-intrinsic innate immunity against pathogenes^28–30^. Therefore, we sought to understand whether the rescue effect could be linked to steady-state cytokine production by expanded Vγ9+Vδ2+ T cell batches, or whether it requires further Vγ9+Vδ2+ T cell activation. We harvested cell supernatants 48-and 72-hours post infection to measure cytokine levels produced by infected wildtype as well as BTN3A pan knockout RKO cells, either with or without co-cultured Vγ9+Vδ2+ T cells. Vγ9+Vδ2+ T cell batches cultured in the presence of IL-2 during expansion as well as within the co-culture assays, released large amounts of cytokines in an uninfected co-culture system. (Figure 3). Lower levels of IL-8, CCL3, CCL4 and GM-CSF were observed in a BTN3A pan knockout co-culture compared to wildtype cells, however, the differences did not reach statistical significance. This indicates a BTN3A-dependent expression and/or release of a specific subset of cytokines by Vγ9+Vδ2+ T cells with very little contribution of epithelial cell cytokines. The very subtle differences are consistent with a low average multiplicity of infection of 1 CFU per cell after 24h under our experimental conditions (Figure S1). Some cytokines showed a trend towards increased secretion upon infection with *L. monocytogenes* (TNF-α, TNF-β, IL-16, CCL3, CCL4, GM-CSF and IFN-γ). We cannot conclude that these non-significant increases play a role in antigen-induced Vγ9+Vδ2+ T cell mediated purging. However, they might reflect activation of a small number or weak activation of Vγ9+Vδ2+ T cells in *L. monocytogenes*-infected cultures with low MOI.

**Figure 3.**
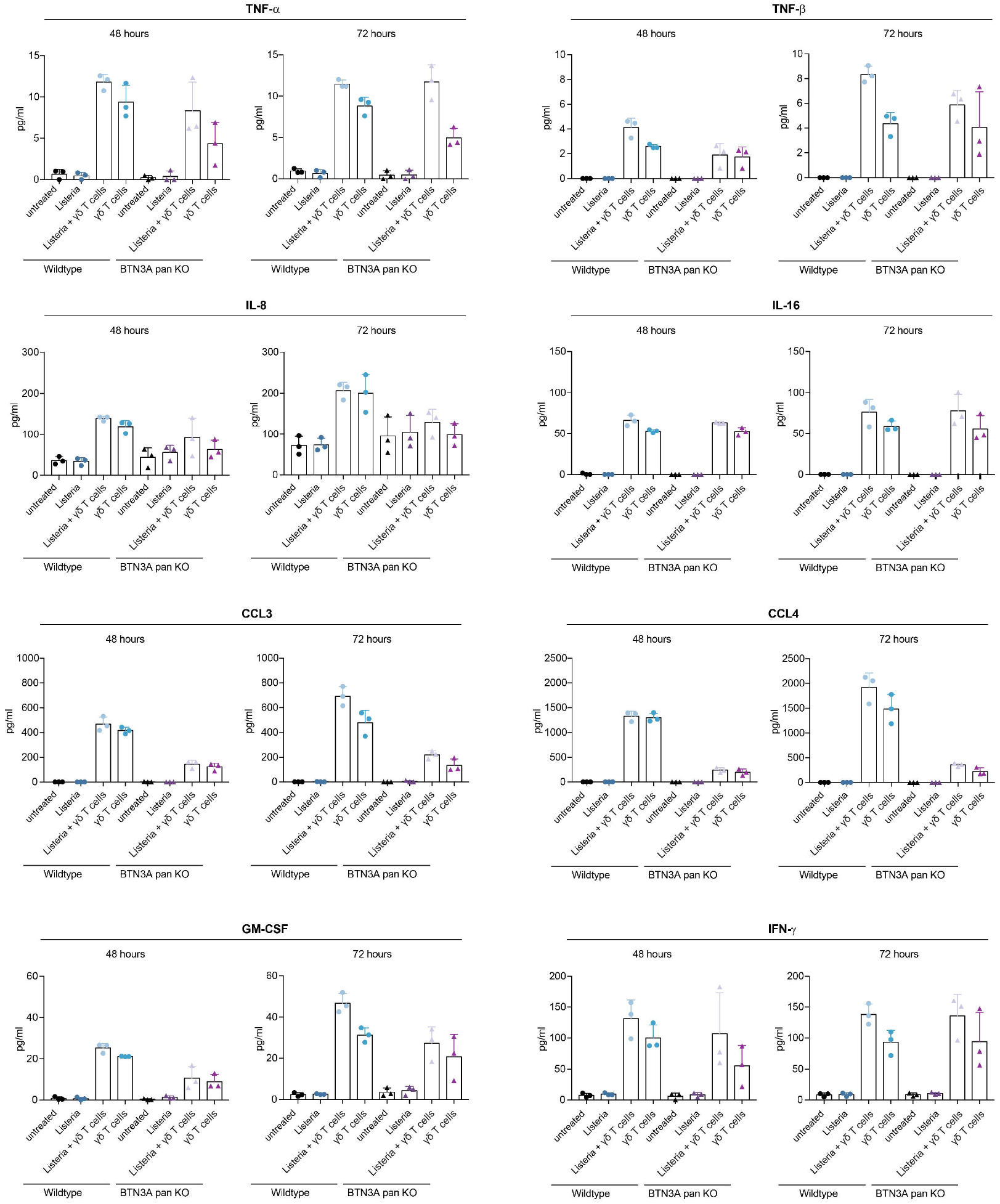
Supernatants of Vγ9+Vδ2+ T cell (donor 1) co-cultures with *L. monocytogenes*-infected or uninfected wildtype and BTN3A pan knockout RKO cells were analyzed using MSD Multi-Spot-Assay. Supernatants were either taken after 48 or 72 hours. Means and standard deviations from three biological replicates are shown.

### 2.4. Clearance of *L. monocytogenes* depends on BTN3A

In contrast to the treatment of epithelial cells with HDMAPP, their infection with *L. monocytogenes* is initially very inefficient and increases gradually via cell-to-cell spreading^31^. We hypothesized that the small number of initially infected target cells were subject to BTN3A-dependent Vγ9+Vδ2+ T cell cytotoxicity. Uninfected cells would escape the cytotoxic activity of Vγ9+Vδ2+ T cells, proliferate, and thus increase the cell index (Figure 4). To monitor the bacterial burden of cells under our coculture conditions, we performed colony-forming unit (CFU) assays in wildtype and BTN3A pan knockout cells infected with *L. monocytogenes* with or without co-cultured Vγ9+Vδ2+ T cells (Figure 5A). Intracellular bacterial numbers were measured every 24 hours over a period of 6 days. While bacterial numbers in the wildtype situation increased immensely over time, they remained at the levels of infection onset in the co-culture. Consistent with the results in Figure 2, this effect was BTN3A-dependent, as the BTN3A pan knockout did not show any difference in bacterial growth when comparing cells infected with *L. monocytogenes* or co-cultured with Vγ9+Vδ2+ T cells. In addition, we performed fluorescence microscopy using a GFP-labeled *L. monocytogenes* strain (Figure 5B). Consistent with the low initial MOI (Figure S1), only a minor fraction of cells was infected with *L. monocytogenes* after 24h, and this fraction was even lower in co-culture with Vγ9+Vδ2+ T cells. After 6 days of infection, *L. monocytogenes* spread throughout the culture, leading to enormous cell stress and cell death. In contrast, the co-cultured cells recovered from the infection, with only very low amounts of intracellular bacteria. Consistent with results in Figure 2, co-culture of BTN3A pan knockout cells with Vγ9+Vδ2+ T cell showed no discernible rescue effect. The epithelial cells exhibited high GFP levels, visible cellular stress, and cell death 6 days after culture alone or together with Vγ9+Vδ2+ T cells.

**Figure 4.**
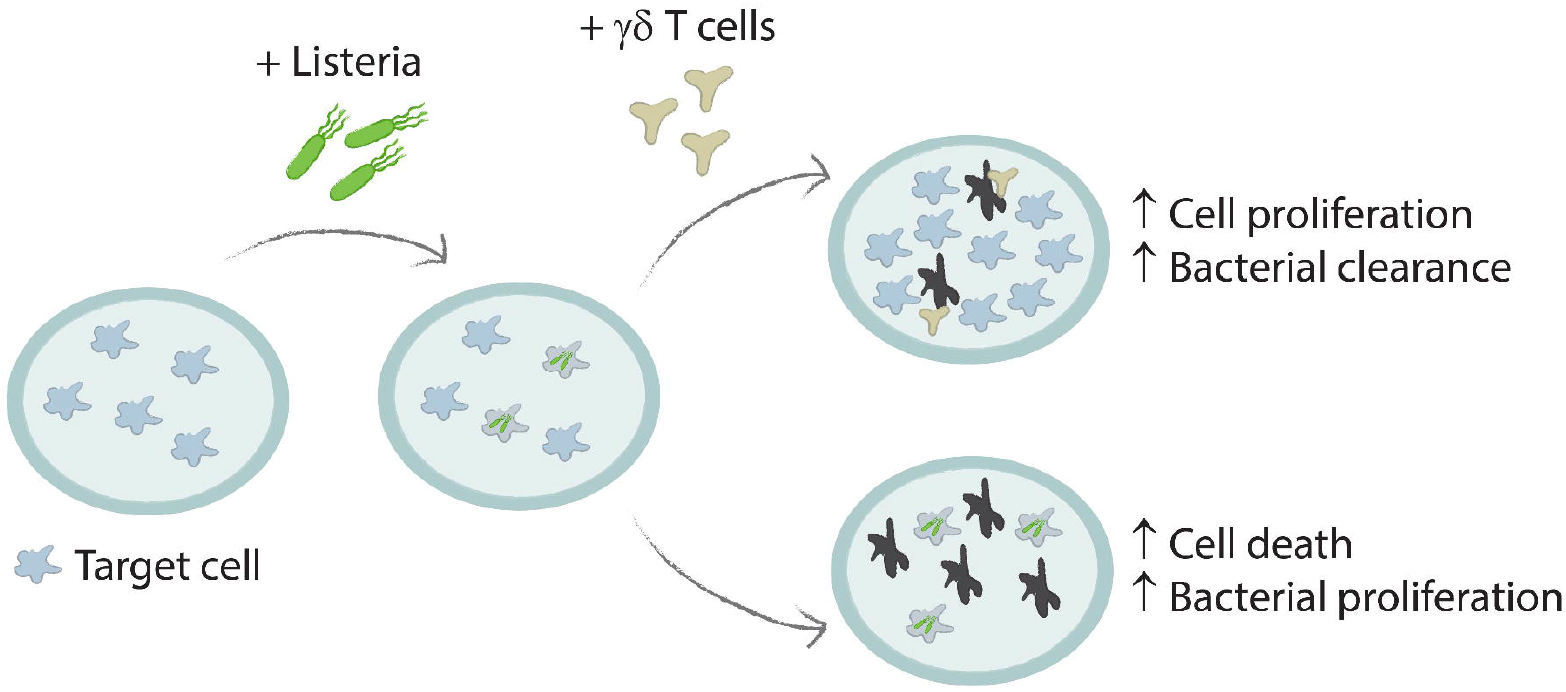
Illustration of L. *monocytogenes*-infected target cells with and without co-culture of Vγ9+Vδ2+ T cells. Upon infection, only a minor amount of target cells takes up L. *monocytogenes*. Over time, L. *monocytogenes* proliferates inside the target cells and spread to the neighboring cells. This results in high amounts of bacteria as well as death of the target cells due to cell-to-cell spreading. In contrast, when co-culturing with Vγ9+Vδ2+ T cells, infected target cells will be immediately attacked by the Vγ9+Vδ2+ T cells reducing the spreading of L. *monocytogenes* to neighboring cells. Thus, infected cells are selectively killed, and uninfected cells will continue growing.

**Figure 5.**
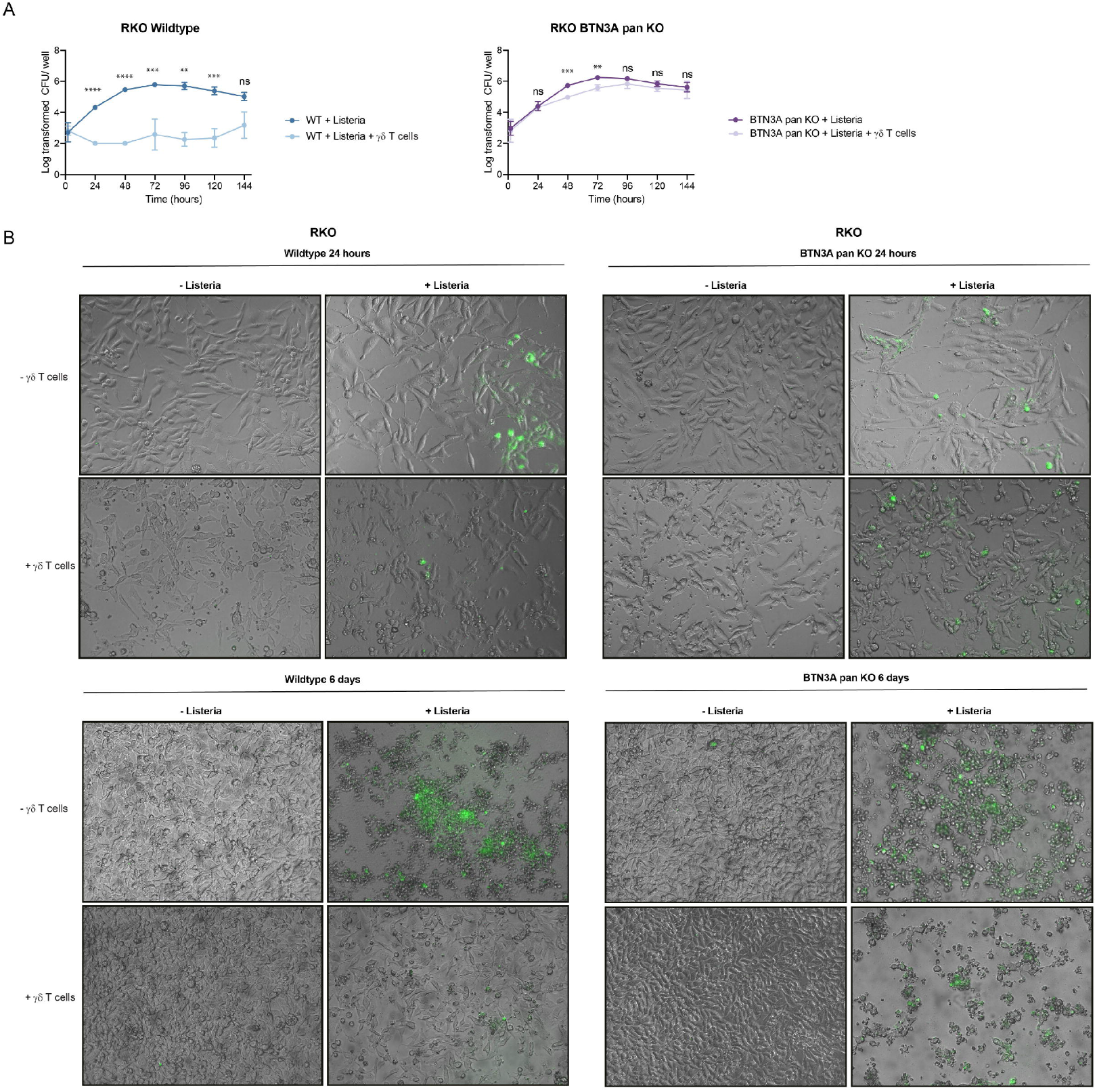
**(A)** Colony forming unit assay was performed using wildtype and BTN3A pan knockout RKO cells infected with *L. monocytogenes* alone or in co-culture with Vγ9+Vδ2+ T cells from donor 1. Cells were lysed every 24 hours over a period of 144 hours and intracellular bacteria were determined by CFU assay. Values represent the log transformed CFU per well of 3 biological replicates. Two-tailed unpaired Student’s t-test was performed using GraphPad Prism comparing L. *monocytogenes* with L. *monocytogenes* + Vγ9+Vδ2+ T cells. P-values: ns P > 0.05; *P ≤ 0.05; **P ≤ 0.01; ***P ≤ 0.001; ****P ≤ 0.0001. (**B**) Wildtype and BTN3A pan knockout RKO cells were infected with GFP-labelled L. *monocytogenes* and/or co-cultured with Vγ9+Vδ2+ T cells from donor 1. Target cells and GFP-labelled bacteria were imaged via microscopy in a live system after either 24 hours or 6 days.

The results are in line with the hypothesis that bacterial clearance occurs at rather early stages of infection by Vγ9+Vδ2+ T cells targeting infected cells. Upon cell killing, uninfected cells start to outgrow and recover from the infection, which is not the case for the BTN3A knockout.

### 2.5. Functional redundancy of BTN3A1 and BTN3A3 regarding the clearance of cell populations from *L. monocytogenes* infection

According to published results, Vγ9+Vδ2+ T cell-induced killing upon HDMAPP stimulation requires the butyrophilin 3A gene BTN3A1 ^20–22,26,27^. We were interested whether the same is true for the killing of cells infected with *L. monocytogenes*. Therefore, we used single knockouts of either BTN3A1 or BTN3A3 (Figure S3) to study their importance during *L. monocytogenes* infection. As our COV362 double knockout of BTN3A1 and BTN3A3 already abolished the ‘rescue effect’, we excluded BTN3A2 as either not involved or functionally redundant for the rescue effect. As expected, a single knockout of BTN3A1 prevented Vγ9+Vδ2+ T cell-induced killing upon HDMAPP stimulation (Figure 6A). In contrast, the knockout of BTN3A3 showed similar results as the wildtype cells (Figure 6B). This behavior of our knockout cells is in agreement with the literature. Surprisingly, the single knockouts of either BTN3A1 or BTN3A3 did not eliminate the ability of Vγ9+Vδ2+ T cells to rescue the cultures infected with *L. monocytogenes*. From these data we conclude that both BTN3A1 and BTN3A3 are functionally redundant in mediating antigen recognition and killing of *L. monocytogenes*-infected target cells by Vγ9+Vδ2+ T cells.

**Figure 6.**
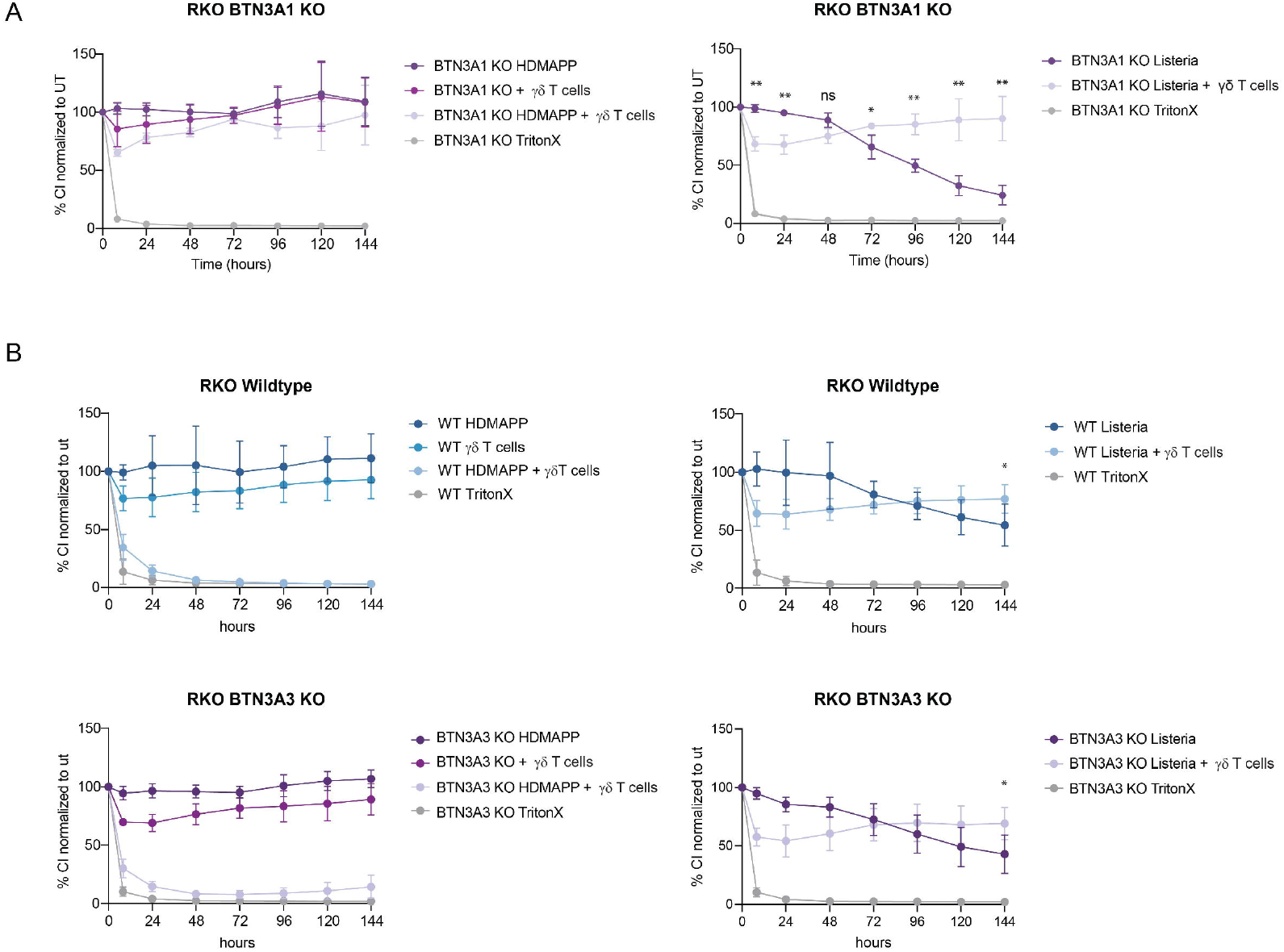
RTCA was performed with BTN3A1 **(A)** and BTN3A3 **(B)** knockout RKO cells as well as wildtype RKO cells (B) either in co-culture with Vγ9+Vδ2+ T cells from donor 1 as well as controls with Vγ9+Vδ2+ T cells alone and cells treated with TritonX. Target cells were either treated with the HDMAPP (left panel) or infected with L. *monocytogenes* (right panel). Cell adherence was monitored over 144 hours. Cell Index of the treatments was normalized to the untreated sample (100%). Two-tailed unpaired Student’s t -test was performed using GraphPad Prism comparing either L. *monocytogenes* with L. *monocytogenes* + Vγ9+Vδ2+ T cells. P-values: ns P > 0.05; *P ≤ 0.05; **P ≤ 0.01; ***P ≤ 0.001; ****P ≤ 0.0001. Mean and STDEV from biological triplicates are shown.

## 3. Discussion

In this study we established an *in vitro* system to investigate responses of Vγ9+Vδ2+ T cells to the infection of epithelial cells with *L. monocytogenes*. With this approach, we advance previous studies mimicking *L. monocytogenes* infection using synthetic phosphoantigens or extracts from heat-killed bacteria. In line with recent literature ^6,12,14,16,32^, we observed that cells exposed to the synthetic HMB-PP analogue HDMAPP were completely eradicated by coculture with Vγ9+Vδ2+ T cells. In contrast, *L. monocytogenes*-infected cultures were protected from complete eradication and showed a “rescue effect” of the culture resulting from the purging of infected cells. Our infection system and the results we obtained differ from those of a recent study with monocyte-derived dendritic cells (Mo-DC) ^9^. Contrasting epithelial cells, Mo-DC resisted the lethal effects of *L. monocytogenes* infection. However, co-culture of infected Mo-DCs with Vγ9+Vδ2+ T cells resulted in their virtually complete elimination by γδ T cell cytotoxicity. Since Mo-DC are capable of phagocytosing bacteria^33^ the different results most likely reflect more efficient infection of phagocytes and a limited potential of noninfected Mo-DCs to expand and rescue the purged culture.

Clearance of infected cells by γδ T cell cytotoxicity requires the activation of Vγ9+Vδ2+ T cells through antigen-activated BTN3A, as indicated by CRISPR/Cas9-mediated KOs of individual, or groups of BTN3A genes in epithelial cells. Investigating supernatants from these co-cultures for their cytokine content, we observed a BNT3A dependency for enhanced TNFβ secretion which correlated with the γδ T cell-mediated rescue of the culture. TNFβ also known as Lymphotoxin A and one of its receptors, LTβR, were reported to be of critical importance for immunity against mycobacterial and *L. monocytogenes* infection in mice ^34,35^. An increased microbial burden was observed in LTβR KO mice in monocytic cells^34^, and also in mice with epithelial cell specific LTβR KO^35,36^. As mice do not have a γδ T cell population equivalent to human Vγ9+Vδ2+ T cells or BTN3A, other cells from other lineages might be the source for TNFβ in the murine system. In our human i*n vitro* infection model, the increased production of TNFβ by with Vγ9+Vδ2+ T cells T cells might be one of the factors increasing the resistance of epithelial cells against *L. monocytogenes*. However, TNFb release cannot account for the cytotoxicity of Vγ9+Vδ2+T cells specifically against the fraction of infected cells in the culture.

Cell cultures lacking all BTN3A family members, or the BTN3A1/3 genes failed to be purged and rescued by Vγ9+Vδ2+ T cells. This rules out an independent role of BTN3A2. Both BTN3A1/3 KO and BTN3Apan KO additionally lack the BTN2A3p, a pseudogene that is transcribed but not further processed and translated. Therefore, a role of BTN2A3p in the presentation of antigens to γδ T cells is highly unlikely. Surprising in the light of recent reports, the rescue effect was observed in both single KO of BTN3A1 and BTN3A3. Contrasting the presentation of the model phosphoantigen HMB-PP, BTN3A1 and BTN3A3 appear to share the ability to present relevant *L. monocytogenes* antigens to Vγ9+Vδ2+ T cells. Alice and colleagues^9^ who reported phosphoantigen-independent Vγ9+Vδ2+ T cell activation by *L. monocytogenes* tested BTN3A dependency using the 103.2 monoclonal antibody. This antibody binds to all ectodomains of the BTN3A family members, and not exclusively to BTN3A1^15^. Its ability to block Vγ9+Vδ2+ T cell activation by *L*. monocytogenes-infected Mo-DC is therefore not contrary to the results we obtained with gene knockouts.

Reportedly, HMB-PP binds to the positively charged pocket of the cytoplasmic B30.2 subunit of BTN3A1, and a single amino acid change in the positively charged pocket of BTN3A3 prevents phosphoantigen reactivity^19^. Therefore, our finding supports the assumption that HMB-PP is not the only antigen inducing the rescue effect in our system. Similar conclusions were drawn by Alice et al.^9^ from results with mutant *L. monocytogenes* strains producing different levels of HMB-PP, but eliciting similar levels of γδ T cell-mediated cytotoxicity against infected Mo-DCs. Overall, the results obtained in experiments with infected cells differ from those generated with heat-killed *L. monocytogenes* extracts and challenge the notion of an exclusive role for HMB-PP as the activating antigen for γδ T cells^6,11^. In this regard, *L. monocytogenes* extracts not subjected to heat treatment were reported to induce increased outgrowth of Vγ9+Vδ2+T cells when compared to heat-treated lysates^8^. This may indicate that heat treatment reduces the potency of HMB-PP, or, alternatively, that it eliminates the activating potential of additional antigens. Moreover, Morita^5^ identified the presence of alkylamine antigens in *L. monocytogenes*^11,37^. Similar to phosphoantigens, alkylamine antigens require cell-cell contact, but not professional antigen presentation via MHC class I, II or CD1 to stimulate Vγ9+Vδ2+T cells. In light of these findings, our observations may indicate an unknown role of BTN3A3 in antigen presentation and the host defense against infections.

## 4. Materials and Methods

### 4.1. Cell Lines

Human colon cancer cell line RKO obtained from the American Type Culture Collection (Cat.No. CRL-2577) were maintained in RPMI 1640 GlutaMAX™ culture medium (Gibco, Cat.No. 61870-010) supplemented with 10% (v/v) heat-inactivated fetal bovine serum (FBS) (Gibco, Cat.No. 26150), 1X Minimum Essential Medium Non-Essential Amino Acids (MEM NEAA; Gibco, Cat.No. 11140-068) and 1X Sodium Pyruvate 100mM (Gibco, Cat.No. 11360-088). Human ovarian cancer cell line COV362 obtained from European Collection of Authenticated Cell Cultures (Cat.No. 07071910) were maintained in DMEM (Sigma, Cat.No. D6429) culture medium supplemented with 10% (v/v) FBS. Lenti-X™ 293T cell line obtained from Clonetech, (Cat.No. 632180) were cultured in DMEM supplemented with 10% Tet System Approved FBS (Takara, Cat.No. 631106). All cell lines were cultivated at 37°C in a humidified atmosphere with 5% CO_2_.

### 4.2. Generation of Primary Vγ9+Vδ2+ T Cell Batches

For the generation of human Vγ9+Vδ2+ T cell batches, buffy coats from healthy donors were purchased from the Austrian Red Cross, who always obtain samples under informed consent in accordance with relevant guidelines, regulations, and internal approvals to ensure ethics and informed consent of donors. PBMCs were isolated from buffy coats by density-gradient centrifugation. In brief, buffy coats were diluted 1:1 in HBSS (Gibco Cat.No. 14170-088) and centrifuged in a Lymphoprep (Axis-Shield, Cat.No. 1114544) containing Leucosep tube (Greiner, Cat.No. 227290) to isolate PBMCs. After 3 washes with HBSS and erythrocyte lysis with ACK lysing buffer (Gibco Cat.No. A10492-01) PBMCs aliquots of 10^8 cells/ml were stored at – 150°C until further use. For Vγ9+Vδ2+ T cell expansion and isolation, PBMCs were thawed and seeded at a cell density of 5*10^6 cells/ml in RPMI 1640 GlutaMAX™ culture medium (Gibco, Cat.No. 61870-010) supplemented with 10% (v/v) heat-inactivated fetal bovine serum (FBS) (Thermo Scientific, #SH30071.03). Cells were treated over night with 0,2 μM HDMAPP*Li (Sigma-Aldrich, Cat.No. 95098) and 125 ng/ml recombinant human IL-2 (Peprotec, Cat.No. AF200-02) and were then expanded over 2 weeks with 125 ng/ml IL-2. At d14, Vγ9+Vδ2 T cells were isolated by negative selection using the γδ T cell isolation kit from StemCell (Cat.No. 19255) according to manufacturer’s protocol. For Donor 11 a δ1TCR antibody cocktail (Cat.No. 18309-A038) was added in addition. Quality and purity of isolated cells was estimated by flow cytometry after surface staining for CD3 (Biolegend, Cat.No. 300428), and γ9 TCR (Biolegend, Cat.No. B235686) or δ2 TCR staining (BD, Cat.No. 0038917) on a FACS CantoII. For both used donors we harvested 99% pure γδ T cells batches, with 88% Vγ9+Vδ2+ content in donor 1 and 91,5% Vγ9+Vδ2+ content in donor 2.

Human primary Vγ9+Vδ2+ T cells from different donors were thawed the day prior the experiment in RPMI 1640 GlutaMAX™ culture medium supplemented with 10% (v/v) heat-inactivated fetal bovine serum (FBS). 10 million cells were resuspended in 10 ml medium and stimulated with 25ng/ml recombinant human interleukin-2 (IL-2) (Peprotech #AF200-02). Vγ9+Vδ2+ T cells were kept overnight at 37°C and 5% CO_2_.

### 4.3. Gene-edited Cell Lines

Human colon carcinoma RKO cell line was stably transduced with pRRL.SFFV-rtTA3-IRES-EcoR-PGK-Hygro and pRRL.TRE3-Cas9-P2A-BFP for tetracyline-induced Cas9 expression. To generate RKO cell lines with a gene knockout for components of the BTN3A family, the following sgRNAs were used:

gControl GGCAGTCGTTCGGTTGATAT

gBTN3Apan AGAACTTCGATTCTGCGGGA

gBTN3A1 CCTGGACGTCTCCTTCTCTG

gBTN3A2 GATGCAGGATACACCCTCCC

gBTN3A3 ATAAAGTGGAGCGACACCAA

Guides were synthetized and cloned into pLenti-sgLib-U6-IT-EF1a-mKate2-P2A-Neo by Genscript. Plasmids were packed into lentiviral particles using the Lenti-X cell line and Lenti-X Packaging Single Shots (VSV-G) (Takara, Cat.No. 631276) according to manufacturer’s protocol. After 48 hours, we harvested viral particle containing supernatants, filtered through a 45 μm SFCA filter and aliquoted and stored them at -80°C until further use. RKO with dox-inducible Cas9 were transduced with the BTN3A guide containing particles and were neomycin selected and tetracyline-induced for 2 weeks to generate polyclonal KO cell batches.

Double knockout clones for BTN3A1 and BTN3A3 were generated by GenScript. Briefly, Cov362 were electroporated with guide containing pSpCas9 (BB)-2A-GFP (PX458), cloned out by single cell dilution and screened for double KO by PCR. The guides used for double KO generation in Cov362 were: gBTN3A1_doubleKO CCTGAGAACTACTAGATGAT gBTN3A3_doubleKO CTGACGTCCCAGTTGTTCCC

## 4.4. Protein Quantification by WES

Seed 2 or 0,33*10^6 cells/well in a 6 well dish and harvest after one or two days respectively. For harvesting, cells were washed with PBS and lysed in 200μl RIPA lysis buffer (EMD Millipore, Cat. No. 20-188) supplemented with cOmplete protease inhibitor cocktail (Roche, Cat. No. 11697498001) and 0,5% n-Dodecyl-β-D-Maltoside (DDM, Avanti Polar Lipids, Cat. No. 850520P). Lysates were collected by scraping, sonicated with 35 mA for 40sec in a water bath and cleared from debris by centrifugation for 10min at 14.000g at 4°C. The protein content of the cleared lysates was estimated using Pierce BCA Protein Assay Kit (Thermo Scientific, Cat.No. 216513) according to manufacturer’s recommendations. BTN3A1 and BTN3A3 protein levels were analyzed using the automated Western Blot System WES from Bio-Techne. In brief, 1μg/μl of cleared cell lysated was loaded onto a ProteinSimple 12-230 kDa 25-Capillary Cartridge (Bio-techne, Cat. No. SM W004), and incubated for 60 min with Antibodies specific to GAPDH (Cell Signaling, #2118, 1:1000), BTN3A1 (Novusbiologicals, Cat. No. NBP1-90750 1:200) or BTN3A3 (Novusbiologicals, Cat.No. 15896-1-AP, 1:200). After 30 minutes incubation with the ProteinSimple Anti-Rabbit Detection Module (Chemiluminescence, Bio-Techne, Cat. No. DM-001), signals were developed using chemiluminescence.

### 4.5. Bacterial Preparation and Infection

*L. monocytogenes* strain LO28 was thawed and grown overnight a day prior the experiment in Brain-Heart Infusion (BHI) medium at 37°C with continuously shaking. After reaching stationary phase, optical density at 600nm (OD_600_) was measured and desired multiplicity of infection (MOI) ratio could be determined. An MOI of 100 was used for experiments with RKO cells, while an MOI of 10 was used for experiments performed with COV362 cells. Bacteria were centrifuged 3 minutes at 10.000 g, washed twice with phosphate-buffered saline (PBS) and resuspended in RPMI 1640 medium supplemented with 10% (v/v) FBS. Bacteria were added dropwise on top of the target cells. After 1 hour of infection, the medium was exchanged to a medium containing a high concentration (50μg/ml) Gentamycin (MP Biomedicals, Santa Ana, US). After a total of 2 hours of L. *monocytogenes* infection, the medium was changed again, and cells kept in a low concentration of Gentamycin (10μg/ml).

### 4.6. Real Time Cell Analysis (RTCA)

The RTCA is a method that allows continuous monitoring of adherent cells. The wells of the associated 96-well E-plates are coated with golden electrodes which monitor the impedance of adherent cells which is further translated into a unit called the Cell Index (CI). The wells of the E-plate were coated beforehand with 50μl Fibronectin (1mg/ml) a day prior the experiment, washed twice with PBS and kept overnight at 4°C. The day of the experiment, the background was measured with 50μl of clean cell medium (3 swipes every 1 minute). Target cells were counted and seeded into the wells of the E-plate. The plate was incubated for 30 minutes at room temperature before being placed back into the monitoring unit which was always kept at 37°C and 5% CO_2_. The interval of the swipes was set to every 5 minutes. Target cells were let to adhere to the wells for 7 hours. Before the treatment, Cell Index of all wells was normalized to 1. At this point, cells were infected with *L. monocytogenes* at the desired MOI (see Section Bacterial Preparation and Infection). After one hour of infection, medium was exchanged to medium containing a high concentration of Gentamycin (50μg/ml). After another hour, cells were either left untreated or treated with 1% (v/v) TritonX-100, HDMAPP (1mg/ml) and/or human primary Vγ9+Vδ2+ T cells with an Effector to Target ratio of 2.5:1. In all conditions, medium was supplemented with a low concentration of Gentamycin as well as recombinant human IL-2 (125ng/ml). The seeding density of the RKO cells line was 2×10^4^ cells/well, whereas we used 1,6×10^4^ cells/well for the COV362 cell line. All treatments were performed in technical triplicates.

### 4.7. MSD Cytokine Analysis

RKO cells were seeded with a density of 2×10^4^ cells/well in a 96-well plate. After 7 hours, cells were infected with *L. monocytogenes* with an MOI of 100. After 1 hour, medium was exchanged to medium containing 50μg/ml Gentamycin. One hour later, Vγ9+Vδ2+ T cells were added to the target cells. Cells were kept in medium supplemented with 10μg/ml Gentamycin and 125ng/ml recombinant human IL-2 at 37°C and 5% CO2. After either 48 or 72 hours, supernatant was harvested. Supernatant was centrifuged 5 minutes at 10.000 g, transferred to a fresh tube and kept at -80°C for 2 weeks to remove any viable bacteria. Afterwards, supernatant was analyzed using the Meso Scale Discovery (MSD) Multi Spot Assay system and the V-PLEX Human Cytokine 30-Plex Kit (MSD, Cat.No. K15054D2) according to manufacturer’s instruction.

### 4.8. Colony Forming Unit Assay (CFU)

RKO cells were seeded with a density of 2×10^4^ cells/well in a 96-well plate. After 7 hours, cells were infected with L. *monocytogenes* with an MOI of 100. After 1 hour, medium was exchanged to medium containing 50μg/ml Gentamycin. One hour later, Vγ9+Vδ2+ T cells were added to the target cells. Cells were kept in medium supplemented with 10μg/ml Gentamycin and 125ng/ml recombinant human IL-2 at 37°C and 5% CO_2_. At the given timepoints, cells were washed twice with PBS and lysed with sterile water. The lysate was further serially diluted in PBS and dilutions were plated on BHI plates. Plates were kept overnight at 37°C and 5% CO_2_ and colonies were counted the next day.

### 4.9. Fluorescent Microscopy

RKO cells were grown on a chambered 8-well microscopy slide (Ibidi, Cat.No. 80807) and kept in phenol red free medium. Target cells were infected for 1 hour with the fluorescent GFP-labelled L. *monocytogenes* strain EGD-cGFP obtained from Olivier Disson, the Institut Pasteur, Unité des Interactions Bactéries-Cellules, Paris F-75015, France^23^. Afterwards, medium was exchanged to medium containing 50μg/ml Gentamycin. After another hour, Vγ9+Vδ2+ T cells were added with medium supplemented with low concentration of Gentamycin (10μg/ml) as well as 125ng/ml recombinant IL-2. Samples were analyzed with a Zeiss Axio Observer Z1 microscope using a 20X objective. Images were further processed with ImageJ software.

## Acknowledgments

We would like to thank all current and previous Boehringer Ingelheim co-workers who contributed to this study by helping with conceptualization, technical expertise, or experimental support; and Johannes Zuber (IMP, Vienna Biocenter) for sharing reagents. The authors gratefully acknowledge a gift of the EGD-cGFP strain from Olivier Disson (Institut Pasteur, Paris, France). Funding was provided by the Austrian Research Fund (FWF) through projects SFB F6106 to TD. KF was supported by the FWF through the doctoral program W1261 Signaling Mechanisms in Cell Homeostasis.

## Author Contributions

KF, VS and TD designed the study and wrote the manuscript. KF, MB and VS performed experiments and analysed the data.

## References

1. Hernandez-Milian, A. & Payeras-Cifre, A. What Is New in Listeriosis? Biomed Res Int 2014, 358051 (2014).

2. Stavru, F., Archambaud, C. & Cossart, P. Cell biology and immunology of Listeria monocytogenes infections: novel insights: Cell biology and immunology of Listeria monocytogenes infections. Immunol Rev 240, 160–184 (2011).

3. Ribet, D. & Cossart, P. How bacterial pathogens colonize their hosts and invade deeper tissues. Microbes Infect 17, 173–183 (2015).

4. Pamer, E. G. Immune responses to Listeria monocytogenes. Nat Rev Immunol 4, 812–823 (2004).

5. Morita, C. T., Mariuzza, R. A. & Brenner, M. B. Antigen recognition by human γδ T cells: pattern recognition by the adaptive immune system. Springer Semin Immunopathol 22, 191–217 (2000).

6. Jouen-Beades, F. et al. In vivo and in vitro activation and expansion of gammadelta T cells during Listeria monocytogenes infection in humans. Infect Immun 65, 4267–4272 (1997).

7. Guo, Y. et al. Human T-cell recognition of Listeria monocytogenes: recognition of listeriolysin O by TcR alpha beta + and TcR gamma delta + T cells. Infect Immun 63, 2288–2294 (1995).

8. Munk, M. E., Elser, C. & Kaufmann, S. H. E. Human γ/δ T-cell response to Listeria monocytogenes protein components in vitro. Immunology 87, 230–235 (1996).

9. Alice, A. F. et al. Listeria monocytogenes-infected human monocytic derived dendritic cells activate Vγ9Vδ2 T cells independently of HMBPP production. Sci Rep-uk 11, 16347 (2021).

10. Ryan-Payseur, B. et al. Multieffector-Functional Immune Responses of HMBPP-Specific Vγ2Vδ2 T Cells in Nonhuman Primates Inoculated with Listeria monocytogenes ΔactA prfA*. J Immunol 189, 1285–1293 (2012).

11. Tanaka, Y. et al. Natural and synthetic non-peptide antigens recognized by human γδ T cells. Nature 375, 155–158 (1995).

12. Begley, M. et al. The interplay between classical and alternative isoprenoid biosynthesis controls γδ T cell bioactivity of Listeria monocytogenes. Febs Lett 561, 99–104 (2004).

13. Frencher, J. T. et al. HMBPP-deficient Listeria mutant immunization alters pulmonary/systemic responses, effector functions, and memory polarization of Vγ2Vδ2 T cells. J Leukocyte Biol 96, 957–967 (2014).

14. Harly, C. et al. Key implication of CD277/butyrophilin-3 (BTN3A) in cellular stress sensing by a major human γδ T-cell subset. Blood 120, 2269–2279 (2012).

15. Palakodeti, A. et al. The Molecular Basis for Modulation of Human Vγ9Vδ2 T Cell Responses by CD277/Butyrophilin-3 (BTN3A)-specific Antibodies*. J Biol Chem 287, 32780–32790 (2012).

16. Wang, H. et al. Butyrophilin 3A1 Plays an Essential Role in Prenyl Pyrophosphate Stimulation of Human Vγ2Vδ2 T Cells. J Immunol 191, 1029–1042 (2013).

17. Vavassori, S. et al. Butyrophilin 3A1 binds phosphorylated antigens and stimulates human γδ T cells. Nat Immunol 14, 908–916 (2013).

18. Rhodes, D. A. et al. Activation of Human γδ T Cells by Cytosolic Interactions of BTN3A1 with Soluble Phosphoantigens and the Cytoskeletal Adaptor Periplakin. J Immunol 194, 2390–2398 (2015).

19. Sandstrom, A. et al. The Intracellular B30.2 Domain of Butyrophilin 3A1 Binds Phosphoantigens to Mediate Activation of Human Vγ9Vδ2 T Cells. Immunity 40, 490–500 (2014).

20. Rigau, M. et al. Butyrophilin 2A1 is essential for phosphoantigen reactivity by γδ T cells. Science 367, eaay5516 (2020).

21. Karunakaran, M. M. et al. Butyrophilin-2A1 Directly Binds Germline-Encoded Regions of the Vγ9Vδ2 TCR and Is Essential for Phosphoantigen Sensing. Immunity 52, 487-498.e6 (2020).

22. Cano, C. E. et al. BTN2A1, an immune checkpoint targeting Vγ9Vδ2 T cell cytotoxicity against malignant cells. Cell Reports 36, 109359 (2021).

23. Balestrino, D. et al. Single-Cell Techniques Using Chromosomally Tagged Fluorescent Bacteria To Study Listeria monocytogenes Infection Processes. Appl Environ Microb 76, 3625–3636 (2010).

24. Lang, F. et al. Early activation of human V gamma 9V delta 2 T cell broad cytotoxicity and TNF production by nonpeptidic mycobacterial ligands. J Immunol Baltim Md 1950 154, 5986–94 (1995).

25. Dang, A. T. et al. NLRC5 promotes transcription of BTN3A1-3 genes and Vγ9Vδ2 T cell-mediated killing. Iscience 24, 101900 (2021).

26. Vantourout, P. et al. Heteromeric interactions regulate butyrophilin (BTN) and BTN-like molecules governing γδ T cell biology. Proc National Acad Sci 115, 1039–1044 (2018).

27. Fichtner, A. S. et al. Alpaca (Vicugna pacos), the first nonprimate species with a phosphoantigen-reactive Vγ9Vδ2 T cell subset. Proc National Acad Sci 117, 6697–6707 (2020).

28. Koga, R. et al. TLR-dependent induction of IFN-beta mediates host defense against Trypanosoma cruzi. J Immunol Baltim Md 1950 177, 7059–66 (2006).

29. Ank, N., West, H. & Paludan, S. R. IFN-lambda: novel antiviral cytokines. J Interf Cytokine Res Official J Int Soc Interf Cytokine Res 26, 373–9 (2006).

30. Ravesloot-Chavez, M. M., Dis, E. V. & Stanley, S. A. The Innate Immune Response to Mycobacterium tuberculosis Infection. Annu Rev Immunol 39, 1–27 (2021).

31. Ortega, F. E., Koslover, E. F. & Theriot, J. A. Listeria monocytogenes cell-to-cell spread in epithelia is heterogeneous and dominated by rare pioneer bacteria. Elife 8, e40032 (2019).

32. Libero, G. D. et al. Selection by two powerful antigens may account for the presence of the major population of human peripheral gamma/delta T cells. J Exp Medicine 173, 1311–1322 (1991).

33. López-Relaño, J. et al. Monocyte-Derived Dendritic Cells Differentiated in the Presence of Lenalidomide Display a Semi-Mature Phenotype, Enhanced Phagocytic Capacity, and Th1 Polarization Capability. Front Immunol 9, 1328 (2018).

34. Ehlers, S. et al. The Lymphotoxin β Receptor Is Critically Involved in Controlling Infections with the Intracellular Pathogens Mycobacterium tuberculosis and Listeria monocytogenes. J Immunol 170, 5210–5218 (2003).

35. Pian, Y. et al. Type 3 Innate Lymphoid Cells Direct Goblet Cell Differentiation via the LT-LTβR Pathway during Listeria Infection. J Immunol Baltim Md 1950 205, 853–863 (2020).

36. Wang, Y. et al. Lymphotoxin Beta Receptor Signaling in Intestinal Epithelial Cells Orchestrates Innate Immune Responses against Mucosal Bacterial Infection. Immunity 32, 403–413 (2010).

37. Bukowski, J. F., Morita, C. T. & Brenner, M. B. Human γδ T Cells Recognize Alkylamines Derived from Microbes, Edible Plants, and Tea Implications for Innate Immunity. Immunity 11, 57–65 (1999).

