## Supplementary Figures for "Vγ9+Vδ2+Tcell-mediated purging of *Listeria monocytogenes*-infected epithelial cells requires butyrophilin 3A genes"

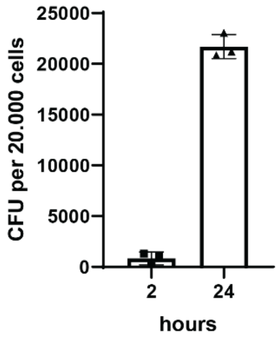


**Supplementary Figure 1**

Colony forming unit assay was performed using *L. monocytogenes* infected wildtype RKO cells. Cells were lysed 2- and 24-hours post infection and intracellular bacterial numbers determined. Values represent the mean of 3 biological replicates.

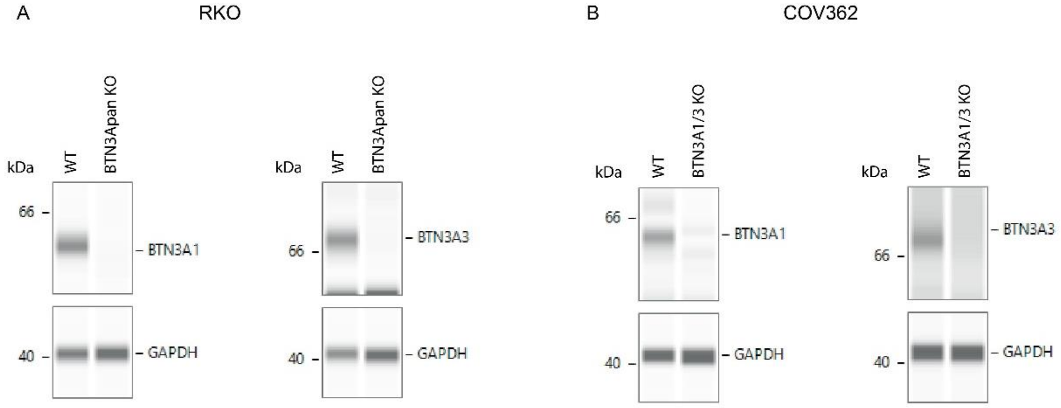


**Supplementary Figure 2**

BTN3A1 and BTN3A3 protein levels of WT RKO cells and RKO cells with knockout of all 3 BTN3A genes (gBTN3Apan). **B)** BTN3A1 and BTN3A3 protein levels of WT COV362 cells and COV362 cells with BTN3A1/3 double knockout (BTN3A1/3 KO) **(B)** using the automated Western blot system WES. One representative WES image is shown **(A, B)**.

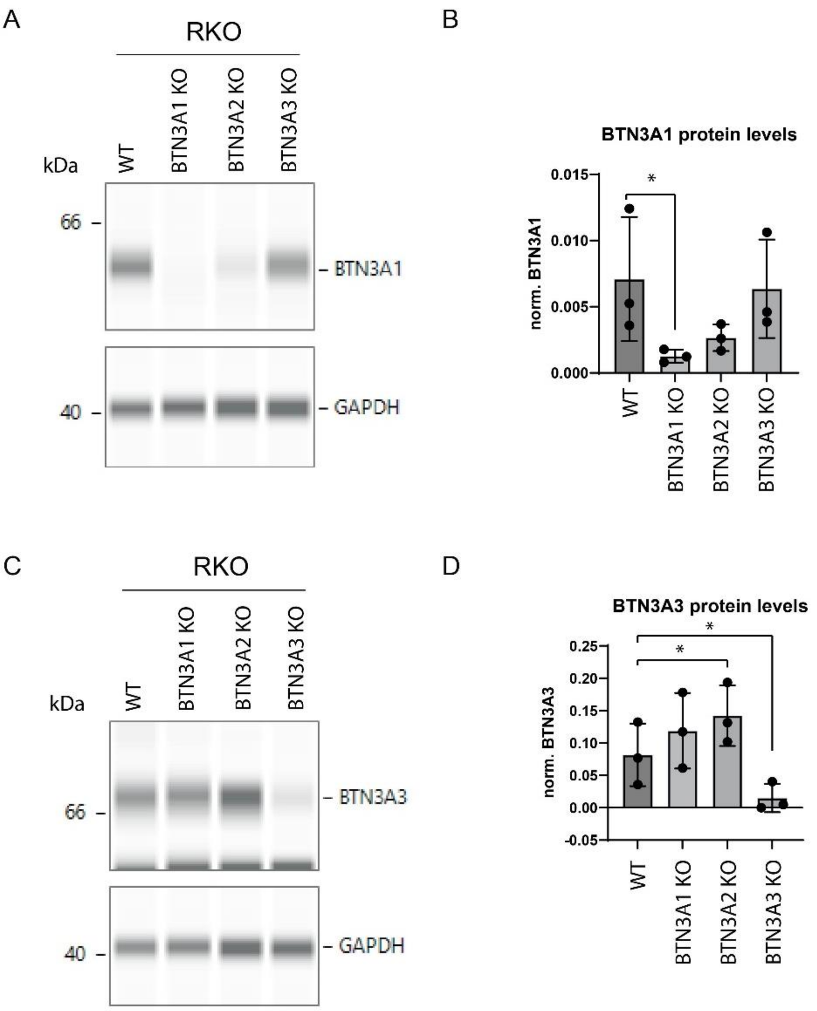


**Supplementary Figure 3**

BTN3A1 and BTN3A3 protein levels from single gene KO of BTN3A family members in RKO cells using the automated Western blot system WES **(A-D)**. For BTN3A1 protein levels one representative WES **(A)** and a summary from 3 biological replicates of relative BTN3A1 intensity values normalized to GAPDH **(B)** are shown. For BTN3A3 also one representative WES **(C)** and a summary of 3 biological replicates of relative BTN3A1 intensity values normalized to GAPDH are shown **(D)**. Mean plus SD are depicted, and statistical significance was tested using GraphPad Prism one-way ANOVA Dunnets multiple comparison test with a single pooled variance; $*P \leq 0.05$ **(B, D)**.
